# Reward invigorates isometric gripping actions

**DOI:** 10.1101/2024.10.25.620324

**Authors:** Rachel M. Marbaker, Ryan C. Schmad, Razan A. Al-Ghamdi, Shruthi Sukumar, Alaa A. Ahmed

## Abstract

Individuals exhibit a propensity to move faster toward more rewarding stimuli. While this phenomenon has been observed in movements, the effect of reward on implicit control of isometric actions, like gripping or grasping, is relatively unknown. How reward-related invigoration generalizes to other effortful actions is an important question. Reward invigorates reaching movements and saccades, supporting the idea that reward pays the additional effort cost of moving faster. Effort in isometric force generation is less understood, so here we ask whether and how reward-related invigoration generalizes to isometric force gripping. And if so, what implicit characteristics of gripping change when there is a prospect of reward? Participants (N=19) gripped a force transducer and the force applied was mapped to radial position of an onscreen cursor. Each trial, a target appeared in one of four locations; increasing grip force moved the cursor toward the target. The gripping action was interchangeable for all target positions. In each block of 100 trials, one target was consistently rewarded, while the other targets were not. When gripping to acquire the rewarded target, participants reacted faster, generated force more rapidly and to a greater extent, while intriguingly maintaining the same accuracy and integral of force over time. These findings support the generalization of reward-related invigoration in isometric force tasks, and that the brain exquisitely trades-off reward and effort costs to obtain reward more rapidly without compromising accuracy or more effort costs than necessary.

**NEW & NOTEWORTHY:** Gripping actions are important for day-to-day tasks, for medical diagnostics like strength and force control, and for choice selection in decision-making experiments. Comparing isometric gripping responses to reward and nonreward cues, we observed reward-based invigoration mediated by selective increases in effort. These findings can be leveraged to provide additional insight into the decision making process, and better understand the effect of reward on movement vigor and the implicit control of accuracy.

## INTRODUCTION

Humans and other animals tend to move physically faster towards rewarding stimuli. The vigor of an action provides a window into the action-selection process. Movements can be completed at an optimal speed with respect to energy efficiency over time; however other factors including end point accuracy, reward acquisition, pain minimization, and time can modulate movement selection and may result in movements with higher energy cost (Berret and Baud-Bovy 2022; van Dieën et al. 2017; Harris and Wolpert 1998; Summerside et al. 2018). For example, for given movement durations, saccadic and arm reaching movements match trajectories that optimize endpoint accuracy (Harris and Wolpert 1998). In these same movements, humans react sooner and move faster to an endpoint that will be rewarded (Summerside et al. 2018; Takikawa et al. 2002). In isometric elbow torque, subjects modulated rate of force development to reduce variability and improve accuracy when hitting target forces (Gordon and Ghez 1987). The importance of time was apparent in a study by Berret and Baud-Bovy (2022) which found that people exerted force to finish a task sooner, even when exerting no force will result in task completion with perfect accuracy in a predetermined duration (Berret and Baud-Bovy 2022). Lastly, reward can not only invigorate movements, but can also, counterintuitively, lead to more accurate movements. These studies demonstrate that action selection is not solely based on metabolic cost and incorporates other sensory and external information in the value function.

The importance of these non-energetic factors is further apparent for action selection in a decision-making context. In a choice between effortful tasks, Korbisch et al. (2022), isolated saccades in choice deliberation from the choice selection saccade and showed that saccade velocities during deliberation increased differentially, with the faster increase in saccade velocities toward the preferred choice (Korbisch et al. 2022). In decision-making studies, it is important to understand not only how choice reflects value, but how the vigor of actions preceding and indicating choice can reveal the subjective value of the selection. For key presses and gripping actions humans respond faster when choice pairs had greater differences in value and when the summed value of choices was greater (Fontanesi et al. 2019; Smith and Peters 2022).

Our goal here is to isolate and understand the effect of reward on the control of isometric gripping actions. Like a movement, gripping incorporates costs such as effort, accuracy, and time. Our understanding of why reward invigorates movement is based on the idea that obtaining reward faster compensates for the metabolic cost of generating a faster movement (Bruening et al. 2024; Shadmehr et al. 2016; Summerside et al. 2024). However, in contrast to movement, the effort of isometric gripping is less understood and it is not clear whether or how reward-based invigoration will generalize to gripping. Historically, metabolic cost has been shown to be proportional to the force-time integral (Crow and Kushmerick 1982; Ortega et al. 2015). The rate of force generation can also be a significant contributor to metabolic cost, beyond the maintenance cost alone (Chasiotis et al. 1987; Hogan et al. 1998; Russ et al. 2002; van der Zee and Kuo 2021). For instance, van der Zee and Kuo (2021) measured metabolic cost during cyclic knee extension against an isometric dynamometer. By subtracting force-time costs and muscle fascicle work from net metabolic cost, they identified a contribution to net metabolic cost that increased linearly with force rate. Taken together, the metabolic cost of gripping is influenced by the magnitude and duration of maintaining force, as well as the rate of force generation. With this study we investigate whether reward will lead to an implicit invigoration of all these determinants, or whether the brain will be selective and optimize for obtaining reward faster without incurring additional metabolic costs.

Adding to the uncertainty surrounding reward-based invigoration of isometric force is that the perception of effort may or may not align with metabolic cost. Perception of the effort of an isometric grip is nonlinear in force and time (Hartmann et al. 2013; Körding et al. 2004; Morel et al. 2017). Thus, it is unclear to what extent objective measures such as force rate, force magnitude and duration influence perception of effort and, by extension, action selection.

If the effort of an isometric force action is not influenced by the rate of force, we would expect no invigorating effect of reward as the effort is independent of the rate at which the force is generated. In contrast, if faster actions are more costly (like force rate), faster high effort actions can be “paid for” by reward. Thus, investigating the effect of reward on gripping can help shed light on representation of effort in isometric tasks. Our experimental design allows for flexible action selection in the isometric force application, and we looked for adjustments in the action related to the anticipation of reward.

Lastly, reward has been shown to augment movement accuracy. When subjects make saccadic eye movements to targets, saccades are both faster and more accurate to targets of higher value (Manohar et al. 2015). This same finding holds in reaching movements for reward (Nikooyan and Ahmed 2015; Summerside et al. 2018). Thus, reward leads to a breakdown of the speed-accuracy tradeoff. One explanation for the speed-accuracy tradeoff is the phenomenon of signal-dependent noise, where larger motor commands result in larger variability (Harris and Wolpert 1998; Jones et al. 2002; Todorov and Jordan 2002). These findings suggest that reward can mitigate the noise that would typically accompany the larger motor commands associated with invigoration. It is unknown whether a similar effect of reward is present in isometric force generation.

Previous studies have shown that the implicit control of movements of our arms and eyes are invigorated by the prospect of higher reward (Summerside et al. 2018; Takikawa et al. 2002). Here, we ask whether the phenomenon of reward-related invigoration generalizes to isometric gripping. We predicted that reward would discount effort and modulate gripping actions. We used various characteristics of gripping action force profiles to explore the nature of effort costs informing action selection.

## MATERIALS AND METHODS

### Subjects

Participants in the gripping experiment, (n = 19, 11 W, 8 M) with an average age of 21.7 ± 0.86 years (mean ± standard error), signed an informed consent form approved by the Institutional Review Board at the University of Colorado. All participants were healthy, with no known neurological disorders or recent injuries affecting the upper limb. All were compensated for their time. Compensation was not contingent on task performance.

### Experimental Setup

Seated participants grasped a force transducer mounted on the handle of an immobilized robotic arm (Shoulder-Elbow Robot; Interactive Motion Technologies; figure 1A). An LCD screen positioned at eye-level displayed a central home circle (radius = 1.25cm), a cursor (0.5cm radius), and the cued target (2cm radius) in one of four positions on any given trial. In this task, the radial position of the onscreen cursor with respect to the home circle was controlled by the isometric gripping force the participant exerted on the robot handle. Each trial started with the cursor in the central home circle, requiring a 500ms pause with gripping force less than 3 N, after which a target marker appeared in one of the four quadrants along the perimeter of a 10cm circle. Using their dominant hand, participants exerted an isometric gripping force on the transducer. As force increased, the cursor moved radially toward the target marker (figure 1B). The position of the target marker at 10cm corresponded to 30N. The cursor position was constrained to the straight-line path in the direction of the target marker. With their gripping force, participants could only control the position along that path. Each target appeared in red. When the target force was achieved, rewarded targets flashed yellow and a pleasing tone was played. Nonrewarded targets turned gray and a neutral tone was played. Gripping force was recorded at 200 Hz and filtered with a fourth order low pass Butterworth with cut off frequency 10Hz.

**Figure 1:**
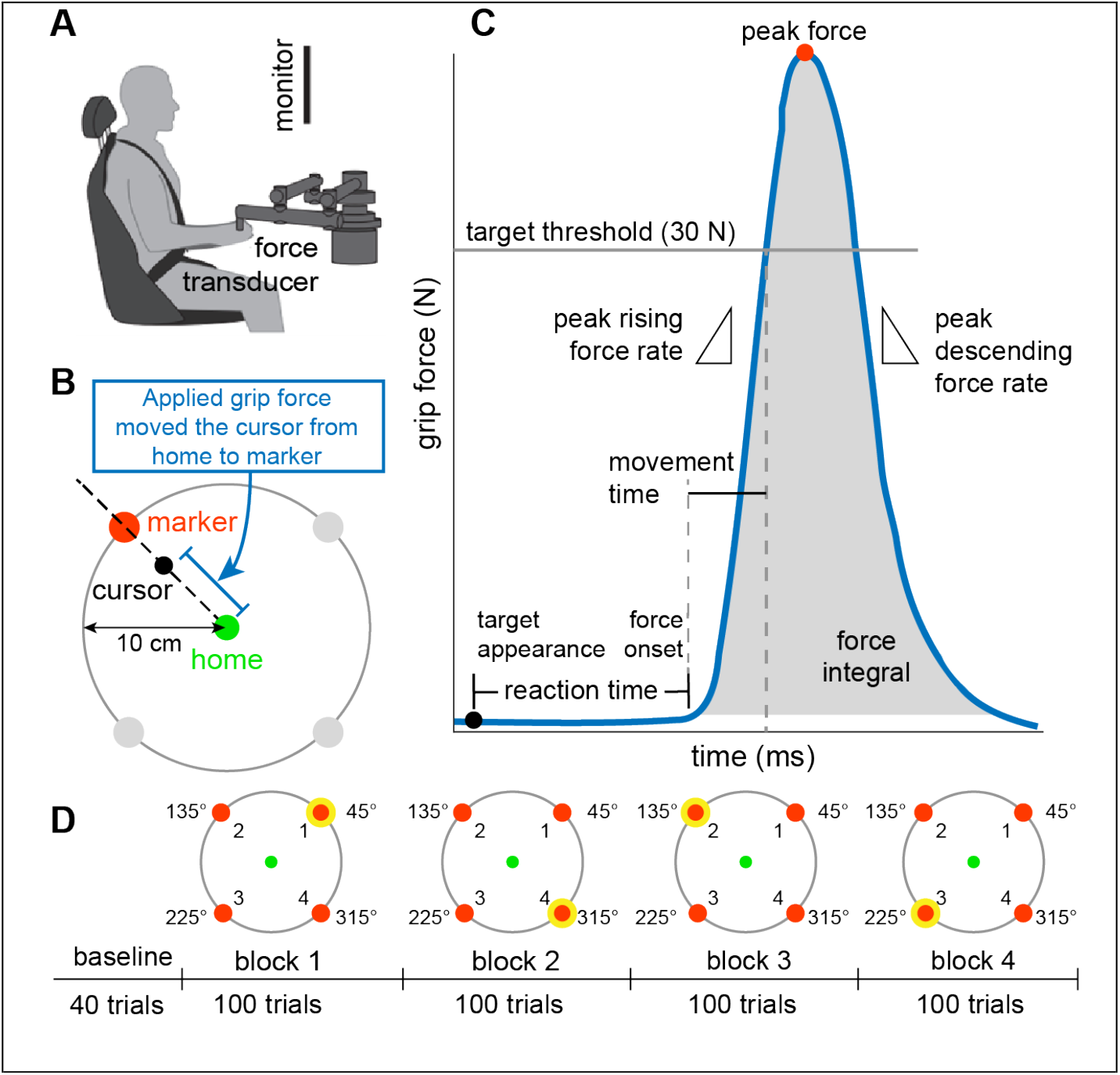
Gripping action task, methods, and metrics. A) Participants were seated, grasping a force transducer mounted on a stationary robot arm. The task was displayed on a screen at eye level. B) Visual grip force feedback. Participants started at low force, with the cursor in the home circle. Once the marker appeared, grip force on the transducer moved the cursor toward the marker. The force to radial distance mapping was linear and the cursor overlapped with the marker at a 30N grip force. C) Gripping action metrics including reaction time, peak rising force rate, movement time, peak force, peak descending force rate, and force integral. D) Sample task structure with familiarization and four blocks of 100 gripping trials. In each of the blocks, one of the markers was rewarded. The order of rewarded markers was randomly assigned for each participant.

To reduce the risk of fatigue or accuracy requirements influencing participant behavior, we set the target force threshold at 30 N, which is low (<10%) with respect to previously reported maximum voluntary contraction forces (353.0 ± 59.72 N for females and 495.2 ± 87.57 N for males (Demura et al. 2003). Since directional movement of the cursor was determined by the target, angular error was always zero.

### Protocol

The first 40 trials contained no rewards and allowed participants to become familiar with the force-to-cursor-position mapping and the task. The next 40 trials contained no rewards and were recorded as baseline gripping. The remaining 400 trials were divided into four blocks of 100 trials each. In each of these blocks, the target in one quadrant was rewarded every time it was cued and the participant achieved the 30N force (Figure 1D). The remaining targets were unrewarded when cued. Over the four blocks the rewarded target was located in each of the four quadrants. Trials within each of the blocks are pseudo randomly assigned in sets of four, such that each set contains a gripping action cued to every quadrant. In each block, 25 of the gripping actions are cued to each marker, so that there are 25 potentially rewarded gripping actions and 75 nonrewarded gripping actions.

### Metrics

The metrics used to characterize and compare the preparation and execution of rewarded and nonrewarded gripping actions we focused on reaction time, peak force, movement time, peak force rate, force integral and time to target (Figure 1C). Reaction time was defined as the time elapsed between target appearance and movement onset. Movement onset was defined as the time where gripping force rate reached 5% of the within-trial peak force rate for a given trial; this is similar to the cursor velocity-based reaction time in Rotella et al (2013). Peak force was the maximum force (>30N) achieved on a trial; because of the force threshold all peak forces exceeded 30 N. Movement time was calculated as the difference between the instant of movement onset and when the force reached the target force of 30N. Peak rising and descending force rate described the highest magnitude rate of force (N/s) preceding and following the peak force for each trial. The starting bound for the force integral used the same bound as reaction time (first time at which the rising rate of force is greater than 5% of the peak rising rate of force). The integral end bound was determined by the opposite: the point at which falling force that is slower than that negative of 5% of the peak rising force rate. Finally, we included a time to target measure, defined as the duration between target appearance and the time that grip force reached the 30 N threshold, reflecting the combined reaction time and movement time.

We removed trials with reaction times exceeding 1s or movement times exceeding 0.5s. Since the gripping action required no directional planning or choice selection, reaction times greater than 1s indicated inattention. Similarly, for movement time there is no accuracy requirement aside from achieving a minimum of 30 N force. Participants were instructed to “quickly grip” the handle and there was no need to sustain the force, so movement times exceeding 0.5 seconds indicated inattention or a misunderstanding of the task. In total, these cutoffs excluded 65 trials or 0.86% of experimental trials. Trials in which the target force was not achieved or required multiple gripping actions to acquire (157 trials or 2.07% of experimental trials) were also removed. Double peak trials were identified manually, verified using MATLAB’s *findpeaks* function and analyzed separately. For descending force rate and force integral, we excluded trials where the force profiles were cut short and did not meet the required boundaries. These, a total of 513 trials, or 5.63%, were excluded only in the calculation of these two metrics.

### Statistics

We sought to determine the effect of reward on the outcome measures. To eliminate any effect of learning related to the changed rewarded cue location or effects of interference from prior reward location, we analyzed only the final 60 trials of each 100-trial block. To account for changing performance over the course of the experiment, differences in reward and nonreward were computed for each 4-trial bin, containing one reward trial and 3 nonreward trials. For each bin, the measures corresponding to the nonreward trials were averaged and compared to the corresponding reward trial using paired t-tests. To calculate temporally-local difference values, the averaged nonreward trial was subtracted from the reward value for each 4-trial bin. Overall differences between reward and nonreward trials for each subject were computed as the averaged bin-differences. We also asked whether reward influenced variability in the outcome measures. Variability metrics were calculated for each subject as the variance of a metric in rewarded and the variance of the same metric in nonrewarded trials. Where variability differences are reported, we subtract the variance of nonrewarded trials from that of rewarded trials. Prior reward effects were assessed using paired t-tests comparing trials preceded by a nonrewarded trial and trials preceded by a rewarded trial for each subject. The effect of target direction was assessed with a linear mixed effects regression model (LMER). To capture the effect of time, we generated LMERs for each force profile metric which included all 400 trials with 100 trials in each block. Trial, normalized by the maximum trial number, was included as a fixed effect to increase the temporal resolution. Reward was included as the additional predictor and a random effect of subject was included for all models. The models were generated for each force profile metric and selected according to marginal *R*^2^ and or AIC if marginal *R*^2^ did not differ.

## RESULTS

In each trial, participants generated an isometric gripping force to move a force-controlled cursor toward one of four possible cued marker locations. One of the marker cues was rewarded and the quadrant of the rewarded cue changed in each of the four, one hundred trial blocks. To complete a trial, participants gripped a transducer to a minimum of 30N; there was no penalty for exceeding the force threshold. We looked at the difference between force profiles of rewarded and unrewarded gripping trials to assess the effect of reward expectation on the preparation (reaction time) and execution (force rate, maximum force, force-time integral) of the gripping action.

### Effects of reward expectation on gripping force

#### Reaction time distributions showed earlier reaction times in the presence of reward

We begin by looking at reaction time, the time it took participants to initiate the gripping action. We computed reaction time distributions (bins = 5 ms) and visualized differences for each participant. Distributions for an example participant (figure 2A) and distributions averaged across participants (figure 2B), reveal that the rewarded distribution was centered earlier than the nonrewarded distribution. Participants initiated force production earlier in the context of expected reward, with shorter reaction times on rewarded trials (RWD-NRWD: -139 ± 32.9 ms; p = 4.96e-04, paired t-test). The difference in rewarded reaction time constituted a 5.14 ± 1.21% reduction from nonrewarded reaction times, suggesting that reward expectation invigorated action preparation (figure 3A).

**Figure 2:**
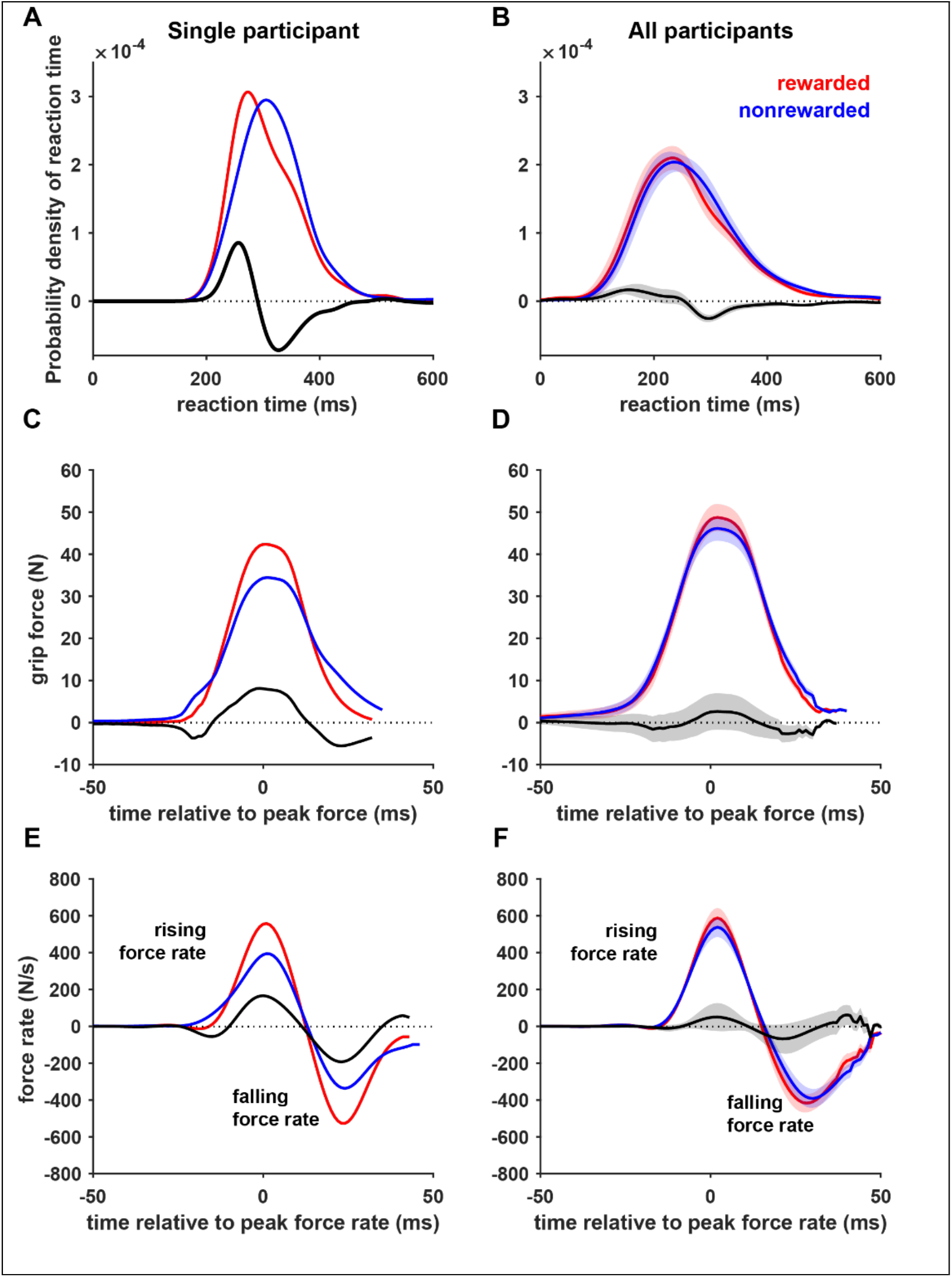
Reward invigorated gripping action. For all plots, data for rewarded trials is shown in red and data for nonrewarded trial is shown in blue. The difference between rewarded and nonrewarded trials (rwd – nrwd) is shown in black. Top: probability density for reaction time. A-B) Rewarded and nonrewarded reaction time distributions for an example participant (A) and across the population (B). The difference between the distributions shows a bias toward earlier reaction times for rewarded trials. C-D) Profiles for the rate of force in during the gripping action aligned to the maximum positive force rate for an example participant (C) and across the population (D) showing higher force rate magnitudes (increasing and decreasing) for rewarded trials. E-F) average grip force profiles aligned to the point of maximum force for an example participant (E) and across the population (F) showing that peak forces tend to be higher for rewarded trials. Plots on the right show mean data for an example participant. Plots on the left show data averaged across participants with standard error shaded.

**Figure 3:**
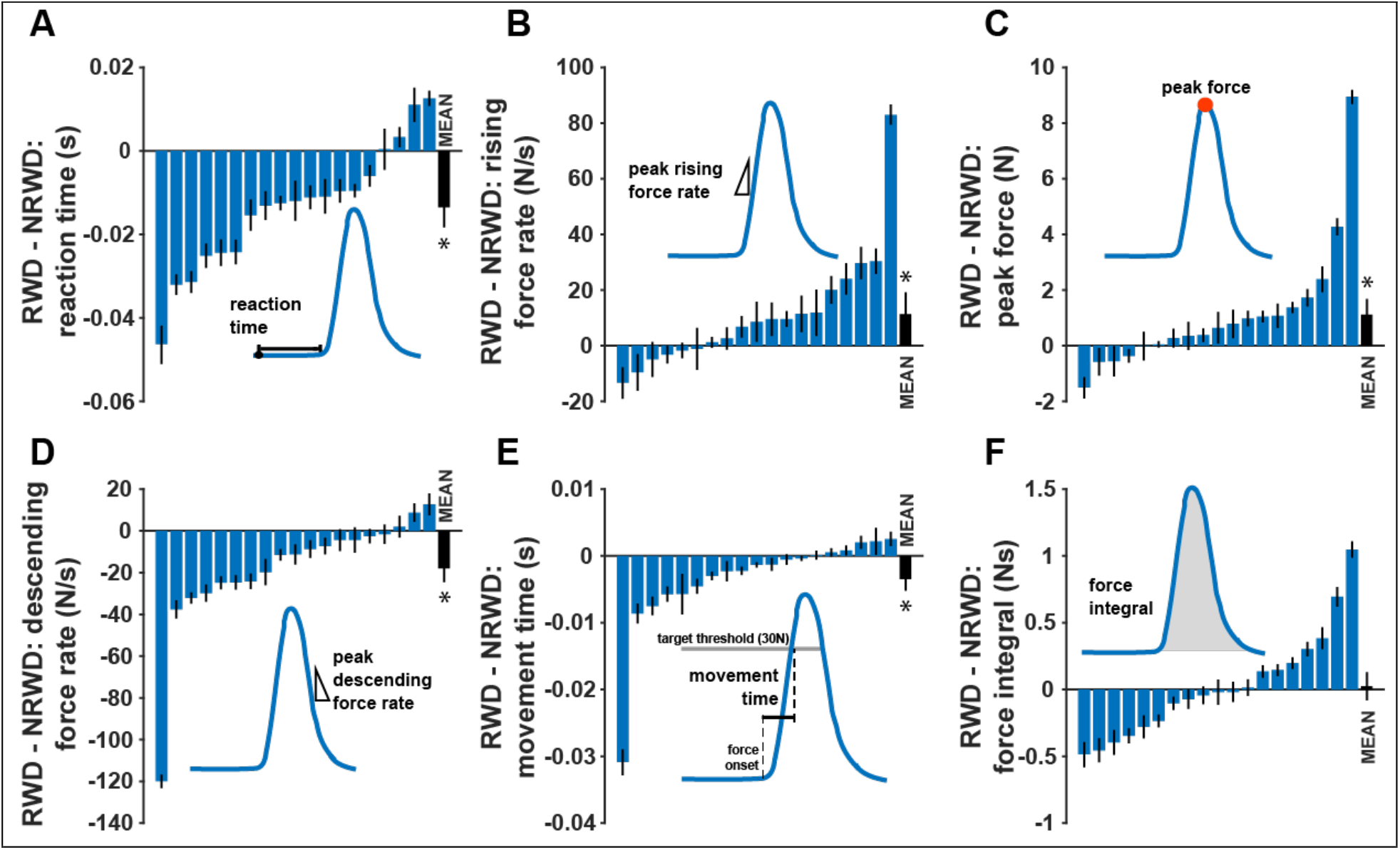
Differences between rewarded and nonrewarded trials for each participant. For each of the barplots, the blue bars represent the mean and standard deviation of RWD – NRWD differences for a participant. The black bar on the right side of each plot shows the mean RWD – NRWD difference across participants with standard error. The average difference between rewarded and nonrewarded trials is shown for each of the grip profile metrics: A) The reaction time difference is negative, indicating longer reaction times for nonreward trials. B) On average the difference in rising force rate is positive, indicating faster force generation for reward trials. C) Peak force has a positive RWD – NRWD difference indicating higher peak forces for reward. D) Because descending force rate is negative, the difference (-RWD – (- NRWD)) indicates faster decreasing force rate for reward trials. E) Like reaction time, the movement time difference is negative, so movement times are shorter for reward trials. F) Finally, the force-time integral did not differ between reward and nonreward trials.

#### Reward expectation increased the rate of force

To examine the execution of the gripping action, we looked at the rate of force as the gripping force increased toward and past the 30 N threshold. We identified the steepest point of the increasing force profile, the peak rising force rate (figure 2C-D). Across subjects, peak rising force rate was higher on trials where participants anticipated reward compared to nonreward (RWD – NRWD: 13.26 ± 5.447 N/s; p = 0.025, paired t-test). This increase in peak rising force rate was equivalent to a 2.4 ± 0.99% of nonreward peak rising force rate. Participants increased their grip force more rapidly in trials with reward expectation (figure 3B).

#### Reward expectation increased peak force

Participant’s gripping force needed to exceed 30N and participants overshot the target force, applying an average of 50.9 ± 2.34 N across all trials (figure 2E-F, positive peaks indicate rising peak force). They applied 1.30 ± 0.541 N (p = 0.027, paired t-test) more force when the target cue was in the reward quadrant (figure 3c). The increase in peak force with reward expectation corresponds to 2.63± 1.093 % of nonreward peak force. Participants grip to a higher peak force in reward trials (figure 3C). Peak rising force rate and peak force on a given trial were positively correlated (R^2^ = 0.641).

#### Reward acquisition increased the magnitude of descending force rate

After achieving and or exceeding the target force, participants reduced the force rapidly to return the cursor to the central home circle and begin the next trial. In trials where the marker was rewarded, descending force rate was faster (more negative) than when the marker was unrewarded.: RWD-NRWD = -22.68 ± 7.560 N/s (p = 0.008, paired t-test). This indicated a more rapid reduction in gripping force after receiving reward (figure 2E - F, negative peaks indicate descending peak force and figure 3D). The magnitude change in descending force rate was 4.44 ± 1.481% of nonreward descending force rate. Peak descending force rate correlated with both peak rising force rate (R^2^ = 0.619) and with peak force (R^2^ = 0.660).

#### Reward expectation reduced movement time

The movement time, the duration between movement onset and reaching the 30N target, was shorter for rewarded trials (RWD – NRWD: -37 ± 1.5 ms, corresponding to a 3.4 ± 1.33% reduction; p = 0.019). Time to target, the duration between target appearance and reaching the 30N target, was also shorter for rewarded trials (RWD -NRWD: -174 ± 42 ms, corresponding to a 4.58 ± 1.11% reduction; p = 6.08e-04). This result confirms that the combination of shorter reaction time and higher peak force rate resulted in faster task completion for faster reward acquisition (figure 3E).

#### Reward did not affect the force-time integral of a gripping action

If we grip an object, both the magnitude of gripping force and the duration of force application contribute to the difficulty of maintaining a grip and the corresponding effort cost of gripping. Force-time integral incorporates both magnitude and duration and may provide an approximation of physical effort expended in each gripping action. The force-time integral of the force profile did not differ between rewarded and nonrewarded trials (RWD – NRWD: 0.038 ± 0.102 Ns, corresponding to 0.46 ± 1.22 % of nonreward force integral; p = 0.7131, paired t-test; figure 3F). No change in force-time integral, despite increased peak force, indicates higher rising and descending force rates may mitigate the increase in peak force. Thus, the combination of increased force rate, increased peak force, and unchanged force integral, suggest an adjustment that maintains the total force-time integral, while acquiring reward more quickly.

#### Across metrics, variance was unchanged for rewarded trial, despite invigoration

We next asked whether reward influenced consistency between trials by comparing the variance of each metric in rewarded trials to the variance of each metric in nonrewarded trials. Despite generating more force and generating it faster, variance remained similar between rewarded and nonrewarded trials for all metrics (reaction time, p = 0.735 (figure 4A); peak risking force rate, p = 0.835 (figure 4B); peak force, p = 0.881 (figure 4C); peak descending force rate, p = 0.893; movement time, p = 0.178; and force-time integral, p = 0.663). Similar variance in the force metrics indicates that invigoration was not accompanied by a reduction in accuracy. Thus, reward expectation may have countered any magnitude-scaled increase in variance.

**Figure 4:**
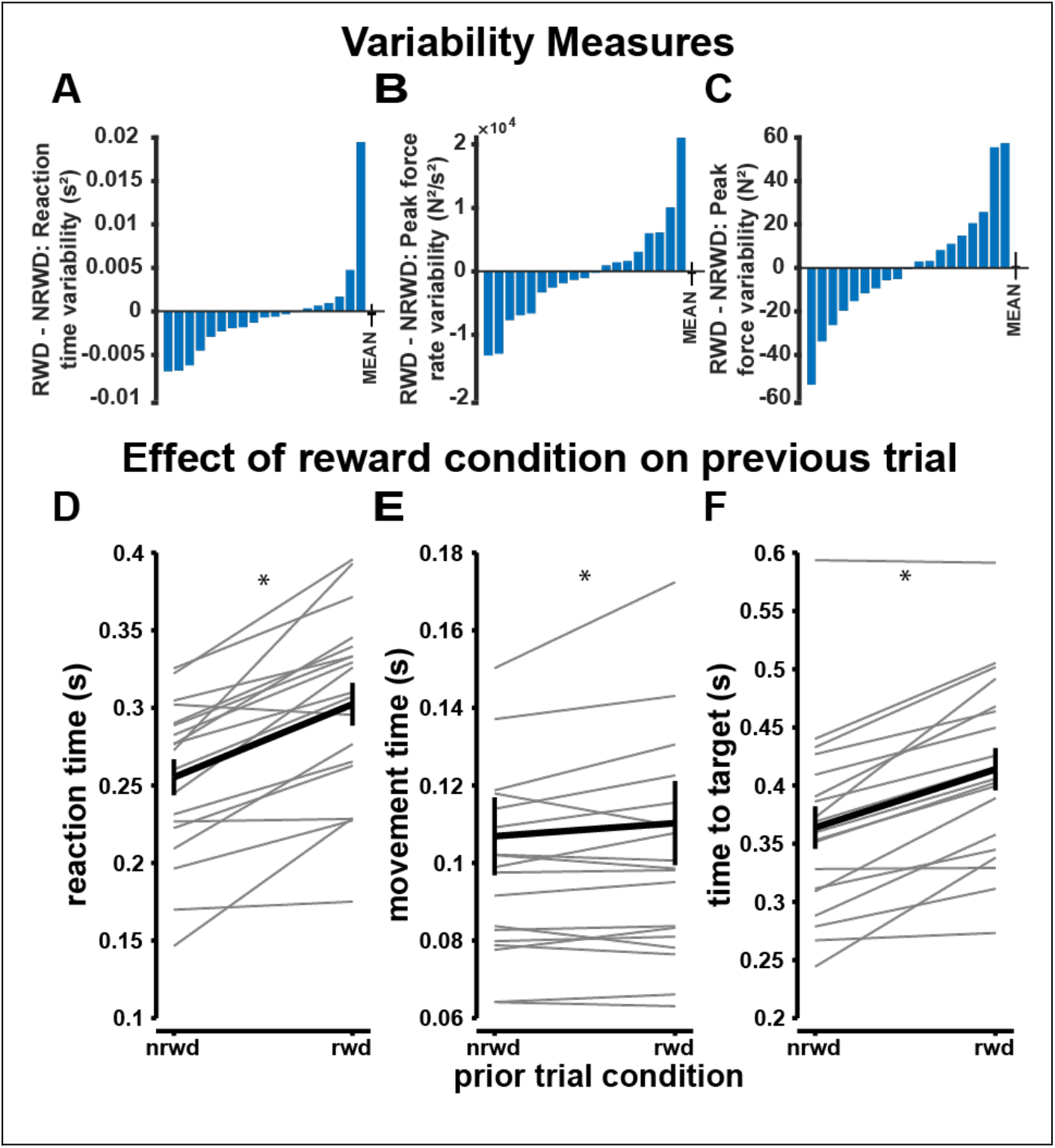
Top: Variability measures. Reaction time (A), peak force rate (B), and peak force (C) were unchanged between rewarded and nonrewarded trials despite reward-related invigoration in all three metrics. Bottom: Effect of Prior Reward. All three time-based metrics, D) reaction time, E) movement time, and F) time to target, were slower when preceded by reward trials than they were when preceded by nonreward trials.

In summary, participants react faster and grip faster with greater force for reward. After reward acquisition, participants release their grip faster. Faster gripping actions do not result in increased variance or in larger force-time integrals, suggesting that the faster rising and descending force rates counter the effect of increased peak forces. If we suppose that vigor generalizes to isometric grip force, reward discounts the effort cost of increasing the force rate and peak force.

### Effects of prior reward condition

If reward influenced the current trial, did these effects persist to the following trial? To investigate the effect of prior reward on gripping actions, we examined differences in the force profiles for trials that had reward or no reward on the previous trial.

Somewhat contrary to our expectation, reaction time, movement time, and time to target were significantly longer on trials with prior reward (reaction time with prior NRWD < reaction time with prior RWD, p = 2.86e-06, figure 4D; movement time with prior NRWD < movement time with prior RWD, p = 0.031, figure 4E; time to target time with prior NRWD < time to target with prior RWD, p = 1.72e-06, figure 4F). The prior reward condition had no effect on peak rising force rate (p = 0.526), peak force (p = 0.253), descending force rate (p = 0.145), or force integral (p = 0.877).

This raised the question of whether statistical reduction in reaction time for rewarded trials was driven by post-reward reaction time slowing. To examine this possibility, we repeated the paired t-tests comparing reaction times for rewarded and nonrewarded trials with trials immediately following reward removed. With post-reward trials removed, reaction time of reward trials remained shorter than that of nonrewarded trials (RWD - NRWD: -8.2 ± 1.18 ms, p = 0.018, paired t-test). Therefore, the invigoration of reaction time on rewarded trials is accentuated, but not driven by slowing in post-reward trials.

### Target location does not change gripping force profiles

In reaching studies, target direction often influences movement speed due to the inertial anisotropy of the arm. In theory, the current experimental paradigm should be insensitive to target direction. To test this, we compared each grip metric with respect to the cued location in the task design. In an LMER analysis that included fixed effects of target direction, trial and reward, the cue location did not generate significant differences in the gripping actions: all estimates for target location factors were nonsignificant, with confidence intervals spanning 0. In Figure 5, the value of each metric, rewarded and nonrewarded is shown for each direction. The nonsignificant coefficients, 97.5% confidence intervals, and p-values for target location for each metric were as follows: reaction time (ms; estimate: 0.68, CI: [-1.171, 2.543], p = 0.470), peak rising force rate (N/s; estimate: -1.988, CI [-4.970, 0.994], p = 0.191), peak force (N; estimate: 0.021, CI: [-0.192, 0.235], p = 0.844), peak descending force rate (N/s; estimate: 0.280, CI: [-2.525, 3.084], p = 0.845), movement time (ms; estimate: 0.38, CI: [-0.339, 1.101], p = 0.300), force integral (Ns; estimate: 0.023, CI: [-0.016,0.062], p = 0.240), and time to target (ms; estimate: 1.05, CI: [-1.027, 3.132], p = 0.321). Since target location was a non-significant predictor in all gripping metrics, we excluded target location as a fixed effect in the following LMER analysis. An RM-ANOVA analysis using a main effect of target location confirmed the null effect of target direction: reaction time: F(3,54) = 0.329, p = 0.805; peak rising force rate: F(3,54) = 0.656, p = 0.579; peak force: F(3,54) = 0.024, p = 0.995; descending force rate: F(3,54) = 0.254, p = 0.859; movement time F(3,54) = 0.212 p = 0.888; and force integral: F(3,54) = 0.223, p = 0.880.

**Figure 5.**
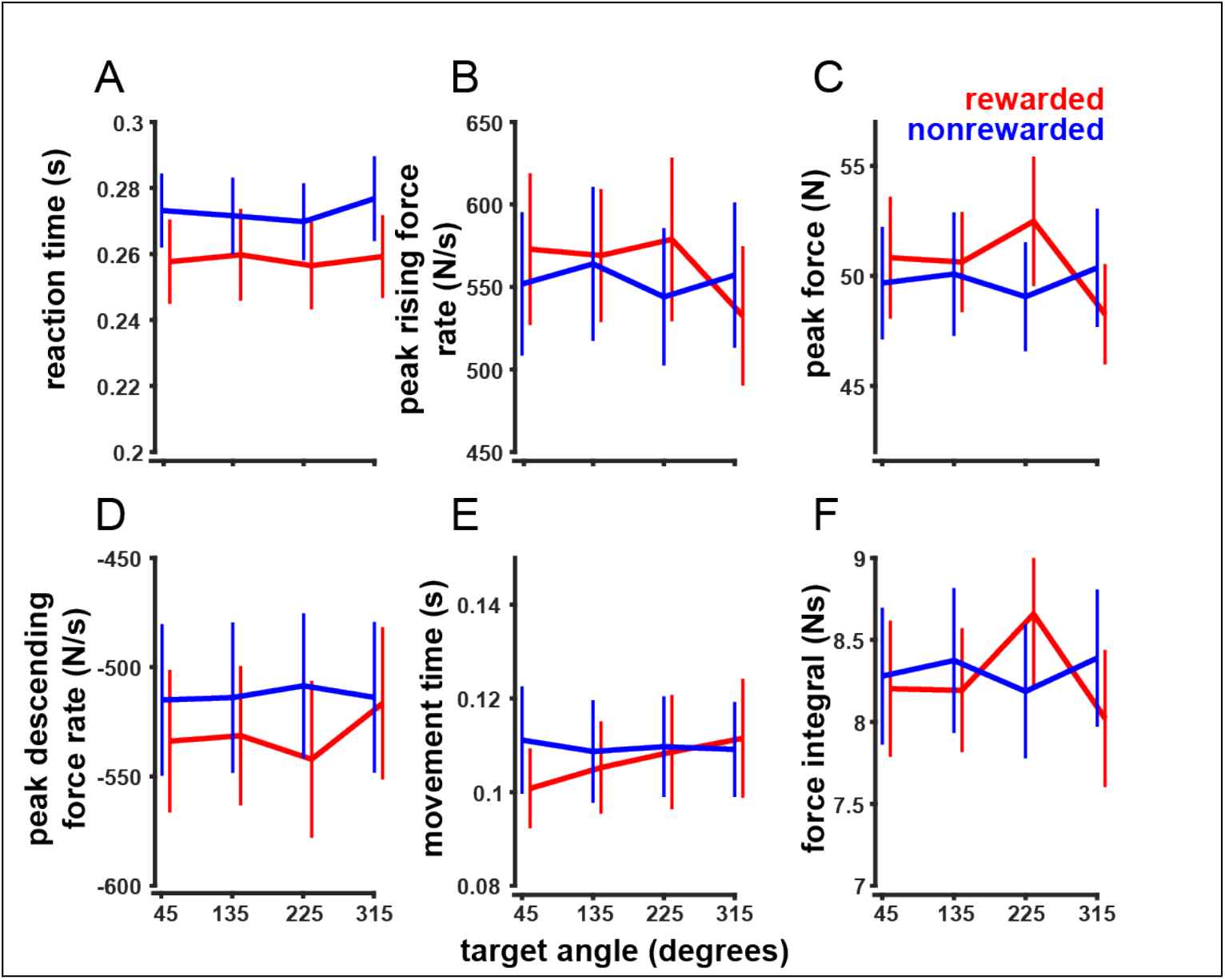
No effect of target direction on grip profile metrics. The direction-specific values for each metric is shown for reward trials (in red) and nonreward trials (in blue) with standard error. There are no directional effects for A) reaction time, B) peak rising force rate, C) peak force, D) peak descending force rate, E) movement time, or F) force time integral.

### Responses changed over time and were sensitive to reward

In our analysis, we noticed that participants appeared to be reacting and gripping faster over the course of the experiment. To more formally capture the effect of time, we generated linear mixed effects regressions (LMER) for each force profile metric (Table 1). All outcome measures, save reaction time, exhibited a significant effect of time. As the experiment progressed, participants generated greater peak force (figure 6C), and both increased and relaxed their grip more quickly (figure 6 B and D). Similarly, both movement time (figure 6E) and time to target decreased with trial number, indicating faster responses. Force integral also decreased with trial number (figure 6F). In contrast, reaction time did not show any slowing or quickening over the course of the experiment (figure 6A).

**Table 1:** Summary of LMER model estimates. The LMER model includes reward and trial with subject was included as a random effect. Coefficients for reward and trial as well as their 97.5% confidence intervals and significance are reported here. (^*^p<0.01, ^**^p<0.001)

|  | Reward estimate | Reward estimate confidence interval | Reward estimate p | Trial estimate | Trial estimate confidence interval (97.5%) | Trial estimate p |
| --- | --- | --- | --- | --- | --- | --- |
| Reaction time (ms) | -13.791 | [-18.576, -9.006] | p = 1.68e -08** | 0.777 | [-6.420,7.974] | p = 0.833 |
| Peak rising force rate (N/s) | 11.186 | [3.488,18.884] | p = 0.004* | 48.100 | [36.523,59.676] | p = 4.48e-16** |
| Peak force (N) | 1.115 | [0.564, 1.667] | p = 1.69e-13** | 1.783 | [0.953, 2.613] | p = 2.57e-05** |
| Peak descending force rate (N/s) | -15.260 | [-22.534, -7.985] | p = 1.11e-11** | -34.790 | [-45.948, -23.630] | p = 1.06e-09** |
| Movement time (ms) | -3.422 | [-5.280, -1.562] | p = 3.12e-04** | -18.779 | [-21.578, --15.979] | p <2e-16** |
| Time to target (ms) | -16.555 | [-21.922, -11.188] | p = 1.56e-09** | -20.801 | [-28.873,-12.730] | p <2e-16** |
| Force-time integral (Ns) | -0.025 | [-0.075, 0.125 ] | p = 0.627 | -0.325 | [-0.475, -0.174] | p = 2.40e-05** |

**Figure 6:**
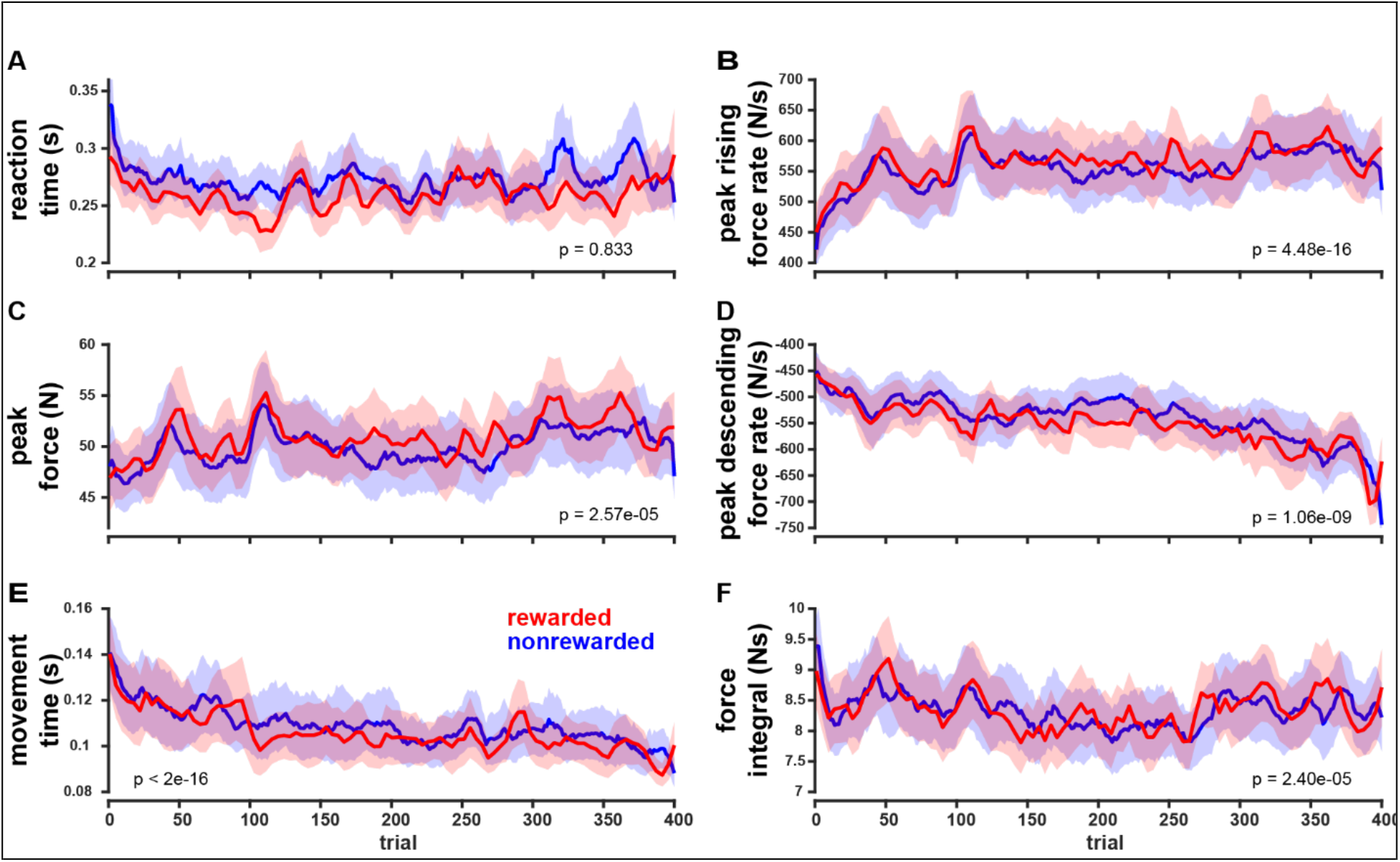
Grip profile metric over the trials of the experiment. Some characteristics of gripping actions changed over the course of the experiment. A) reaction time was unchanged over trials. B) peak rising force rate increased over the trials of the experiment, as did C) peak force. D) Peak descending force rate increased in magnitude (becoming more negative) across trials. Unlike reaction time, movement time (E) decreased over the experiment. Finally, force integral (F) decreased very slightly across experimental trials. Data from reward trials (shown in red with shaded standard error) was smoothed using a moving average with a span of three (3). Data from nonreward trials (shown in blue with shaded standard error) was smoothed using a moving average with a span of nine (9) since there are three times as many nonreward trials as reward trials.

The effect of reward in the LMER was consistent with findings from our earlier t-test analysis: in summary: rewarded trials showed decreased reaction time, movement time, and time to target, showed increased rising force rate, peak force, and descending force rate magnitude, and showed no reward-related change to force-time integral.

### Reward did not reduce double peak failure rate

If participants failed to reach the 30N threshold with the first gripping action in response to the target cue, they gripped again, producing a force profile with two peaks. These trials were excluded from the previous analysis but may provide insight related to risk or sensitivity to reward. Takikawa et al. (2002) observed reduced numbers of incorrect saccades in reward trials. Are participants more likely to undershoot the threshold, creating a double peak, on nonrewarded trials?

Participants had as few as zero and as many as 27 total double peak trials (corresponding to 0% and 11.25% of of a subject’s experimental trials). For each subject, we tallied the double grip actions for rewarded and nonrewarded trials. Noting that there are three times as many nonreward trials represented, we generated expected values for the number of double peaks. As count data, differences in double peak occurrences from the expected values were examined for each subject using chi-squared tests. For all subjects, the occurrences of rewarded double peak trials and nonrewarded double peak trials were similar. This finding held when instances of double peaks were summed across subjects (total of 293 trials), indicating that participants were equally likely to undershoot the force threshold for reward and nonreward trials (chi-squared test, p = 0.1815).

## DISCUSSION

Isometric gripping actions for future reward were characterized by shorter reaction times, faster rising force rate, higher peak force, and faster falling force rate. Rewarded and nonrewarded trials had similar reaction time and peak force variability and the number of double peak trials, indicating that shorter reaction times and higher rising force rates for reward did not result in greater variability with respect to peak force or task success. Further, rewarded and unrewarded trials did not differ in force-time integral, a determinant of metabolic cost in isometric force tasks. Overall, this suggests that participants selectively increased effort to acquire reward faster and more accurately, yet mitigated the increase in effort by conserving the force-time integral.

Reaction time was significantly shorter for rewarded trials than for unrewarded trials. This is consistent with findings in reaching and saccades (Manohar et al. 2015; Sedaghat-Nejad and Shadmehr 2021; Summerside et al. 2018; Takikawa et al. 2002; Yoon et al. 2019). Peak rising force rate increased for reward trials, akin to the greater peak velocity with reward observed in reaching and saccades. Together with shorter reaction time, faster rising force rate suggested a strategy to acquire reward sooner. Indeed, movement time was faster for rewarded trials. Overall, our results in isometric gripping parallel the main effects of reward observed in movement of the eyes and the arm: participants reduce time to reward acquisition via a combined strategy of shorter reaction time and faster rate of force.

Rewarded trials had a higher peak force. This is similar to the effect of reward observed on reaching movements, where participants overshoot rewarding targets more than they overshoot neutral targets (Summerside et al. 2018). Given inherent motor noise, this overshoot may reflect a desire to increase the probability of reward by shifting the distribution of potential gripping actions higher above the target (Harris and Wolpert 1998; Jones et al. 2002). In our findings, peak force variability was unaffected by reward, so by increasing peak force, participants are increasing the likelihood that the force will exceed the threshold, resulting in reward. Thus, it is plausible that increasing the peak force on rewarded trials reduced the risk of missing out on the reward. Prior work on isometric elbow torque control, noted participants’ tendency to undershoot force targets when instructed to be “accurate” and their tendency to overshoot force targets when instructed to be “fast” (Gordon and Ghez 1987). In our study, participants may be taking advantage of the minimum-only force threshold to acquire reward faster without force rate slowing since there are no overshoot consequences.

Interestingly, we found no difference in force-time integrals between rewarded and unrewarded grip trials. This indicates that participants could increase rising peak force rate and peak force while maintaining the force integral. In this way, they acquired reward more quickly and without increasing the impulse of the action, possibly mitigating the associated increase in effort.

A cornerstone of current theories of movement control is the role of signal-dependent noise (Harris and Wolpert 1998; Jones et al. 2002). Greater force is indelibly accompanied by greater variance. We do not see this effect in our results. Participants generate greater force for reward, but not at the expense of greater variability. Similar results have also been observed in reaching and saccades, where higher vigor does not reduce endpoint accuracy (Manohar et al. 2015; Summerside et al. 2018). It will be interesting for future investigations to understand the basis of such reward-related implicit improvements in motor control.

Why should reward invigorate action? The rationale behind reward-based invigoration is anchored in the presumption that there exists a tradeoff between acquiring reward sooner and exerting additional effort to act faster. Slower actions take longer and delay reward, and faster actions are more effortful, but acquire reward sooner. The metabolic cost of isometric force generation is related to both the force-time integral as well as the rate of generating that force (Hogan et al. 1998; Russ et al. 2002; van der Zee and Kuo 2021). However, it is not clear whether the perception of effort in an isometric task involves solely the force-time integral or also incorporates cost information related to the rate of force generation. In our task, these two effort formulations—either including or excluding rate-related costs—make different predictions. If, in the model, effort is solely the cost of maintenance (i.e. the force-time integral), then the rate of generating force and acquiring the target faster has no effect on effort. Thus, the expectation is that reward will not invigorate movement. In contrast, if effort depends on the rate of force generation, then similar to the effects of reward on reaching, reward will pay the cost of faster isometric force generation. Thus, our findings that reward invigorated gripping actions further implicate force rate as an important contribution to effort cost.

Our gripping action task was free from mechanical directional effects, but force was mapped to radial cursor position and participants may have perceived the task as similar to a reaching movement. In planar reaching, individuals prefer to reach along the axis of movement that minimizes the inertia of the arm (Cos et al. 2011; Goble et al. 2007; Schweighofer et al. 2015). Individuals also move more quickly in low inertia reach directions, where the arm has lower effective mass (Gordon et al. 1994). Directional effects in our data would indicate the generalization of a location-based perception of effort. The absence of directional effects in gripping actions verifies that mechanically based directional effects in reaching are not transferred to other modalities based on spatial cues from the visual feedback. This information may be of value for researchers interpreting isometric force data or using location-based cues in other paradigms.

Further understanding the dynamics of gripping in response to reward is important as it may shed light on whether the brain globally invigorates all actions in pursuit of reward. In a broad interpretation, behavior we observed in isometric gripping was similar to findings in reaching and saccades: the presence of reward reduced reaction time, increased the rate of the action toward the goal, and increased the magnitude of the action. Our findings suggest a generalization of reward-related invigoration beyond spatial movements to isometric force. Gripping tasks are advantageous in the study of effort because, they are free from the directional effects intrinsic to reaching studies wherein movements to different target locations have different effort cost due to interaction torques and inertial resistance (Goble et al. 2007; Gordon et al. 1994; Schweighofer et al. 2015).

We also observed faster, higher force gripping actions as the trial progressed. There was no change in reaction time, but changes in rising force rate (increase), peak force (increase), descending force rate (magnitude increase), movement time (decrease), and time to target (decrease) were consistent with invigorated gripping actions related to the increasing trial number (regardless of the reward or nonreward condition of the stimulus). This gradually invigorated behavior is consistent with studies on reaching, where participants reached faster as the experiment progressed, but opposed studies of saccades which show slowing over the course of an experiment (Summerside et al. 2018; Xu-Wilson et al. 2009).

Gripping, like reaching and saccades, can also offer a window into human decision-making when it is used as a means of choice selection in decision-making paradigms. Smith and Peters (2022) examined gripping dynamics in the context of a choice-task between a low, immediately available reward and a high, delayed reward. They found peak force (amplitude) was higher in choice trials where the immediately available option was higher. No differences related to the immediately available reward were detected for the centroid and width. Centroid and width are indirectly related to reaction time and force rate, but differences in the task design make the metrics difficult to compare. Smith and Peters’ task had an unlimited response time and the choice difficulty changed across trials. Furthermore, it is not clear that findings in a two-choice paradigm, which incorporate deliberation and selection, generalize to a forced-choice. For example, Korbisch et al. (2022) looked at saccade vigor in a choice deliberation and choice selection where deliberation and selection were separated by a waiting period. While saccades during deliberation increased in velocity toward the preferred option, during choice selection when the decision was already made, there was no difference in saccade velocity (Korbisch et al. 2022). Thus, invigoration in a choice task likely represents mingled effects of reward and deliberation, whereas invigoration in the forced-choice paradigm can be more cleanly attributed to reward.

## Limitations

Though our results show no difference in gripping actions related to target direction and invigoration for rewarded targets, our study design requires no differentiation of actions toward a particular target, and thus could be completed without using the visual information presented. Additionally, our study used very simple reward feedback, a flash, pleasing tone, and the addition of four points. These two factors may have unintentionally reduced participants’ engagement, especially as the experiment progressed. Further, the primary reward feedback (flash and tone) was nonquantitative, and we did not modulate reward magnitude. Neither points nor any other metric of performance in the task were linked to compensation.

Response times were limited, but the intertrial duration was not controlled. To start a trial, participants were required to reduce their grip force below 0.4 N for 500 ms before the next target cue appeared. The remaining time in each trial depended on how quickly the participant produced the required 30N force and returned to below the 0.4N threshold. Accordingly, participants with faster reaction times and faster gripping actions could move through trials more rapidly, resulting in a higher average reward rate. Reward rate and history has been shown to modulate saccades, reaching movements, and gripping, so differences between subjects’ action times may have been exacerbated by reward rate effects (Shadmehr and Ahmed 2020; Sukumar et al. 2022; Yoon et al. 2018). In our data, we also noted significantly longer reaction times in trials immediately after reward. This post-reward slowing has been observed in saccades and Takikawa et al. (2002) proposed potential mechanisms that included post-reward satiety and decreased motivation due to the low probability of a second reward immediately following a first (Takikawa et al. 2002). Our trial pseudorandomization relied upon four-trial sets, so that the median number of repeat rewards was 8. In the full data set, the probability of repeated reward was 0.029. Post-reward slowing could be related to decreased motivation; however, since participants’ rapid progress through trials, the visual and auditory feedback for reward may have reduced subject’s ability to respond to subsequent cues.

## Conclusion

Humans performing an isometric gripping action exhibited shorter reaction times, generated force more rapidly, and reached greater force magnitude when they expected reward. Despite the increase in peak force, accuracy and the integral of force maintained over time were unchanged. These findings support the generalization of reward-related invigoration in isometric force tasks, and that the brain exquisitely trades-off reward and effort costs to obtain reward more rapidly without compromising accuracy or more effort costs than necessary.

## Notes

**GRANTS:** This work was supported by grants from the National Institutes of Health (1R01NS096083) and the National Science Foundation (CAREER award 1352632) to AAA and GRFP (DGE 2040434) to RMM)

**DISCLOSURES:** The authors report no potential conflicts of interest.

### Competing Interest Statement

The authors have declared no competing interest.

